# RhoA activation promotes glucose uptake to elevate proliferation in MAPK inhibitor resistant melanoma cells

**DOI:** 10.1101/2024.01.09.574940

**Authors:** Vasanth Siruvallur Murali, Divya Rajendran, Tadamoto Isogai, Ralph J. DeBerardinis, Gaudenz Danuser

## Abstract

Cutaneous melanomas harboring a B-Raf^V600E^ mutation are treated with immune check point inhibitors or kinase inhibitor combination therapies relying on MAPK inhibitors (MAPKi) Dabrafenib and Trametinib (Curti and Faries, 2021). However, cells become resistant to treatments over the timespan of a few months. Resistance to MAPKi has been associated with adoption of an aggressive amoeboid phenotype characterized by elevated RhoA signaling, enhanced contractility and thick cortical filamentous actin (F-actin) structures (Kim et al., 2016; Misek et al., 2020). Targeting active RhoA through Rho-kinase (ROCK) inhibitors, either alone or in combination with immunotherapies, reverts MAPKi-resistance (Misek et al., 2020; Orgaz et al., 2020). Yet, the mechanisms for this behavior remain largely unknown. Given our recent findings of cytoskeleton’s role in cancer cell proliferation (Mohan et al., 2019), survival (Weems et al., 2023), and metabolism (Park et al., 2020), we explored possibilities by which RhoA-driven changes in cytoskeleton structure may confer resistance. We confirmed elevated activation of RhoA in a panel of MAPKi-resistant melanoma cell lines, leading to a marked increase in the presence of contractile F-actin bundles. Moreover, these cells had increased glucose uptake and glycolysis, a phenotype disrupted by pharmacological perturbation of ROCK. However, glycolysis was unaffected by disruption of F-actin bundles, indicating that glycolytic stimulation in MAPKi-resistant melanoma is independent of F-actin organization. Instead, our findings highlight a mechanism in which elevated RhoA signaling activates ROCK, leading to the activation of insulin receptor substrate 1 (IRS1) and P85 of the PI3K pathway, which promotes cell surface expression of GLUT1 and elevated glucose uptake. Application of ROCK inhibitor GSK269962A results in reduced glucose uptake and glycolysis, thus impeding cell proliferation. Our study adds a mechanism to the proposed use of ROCK inhibitors for long-term treatments on MAPKi-resistant melanomas.

## Introduction

Cutaneous melanomas can be classified into four subgroups based on their acquired mutation in either *BRAF* (50%), the small GTPase *NRAS* (25%), or the tumor suppressor and negative regulator of Ras, neurofimbrin 1 (*NF1*) (14%), and triple negative melanomas (Curtin et al., 2005; Czarnecka et al., 2020; Hayward et al., 2017). The V600-to-E gain-of-function mutation in B-Raf (B-Raf^V600E^) constitutively activates the mitogen activated protein kinase (MAPK) pathway (Hodis et al., 2012) independently of upstream regulators and stimuli, and promotes proliferation, survival and tumor progression (Davies et al., 2002; Gray-Schopfer et al., 2007). B-Raf^V600E^ served as one of the first candidates for the development of targeted therapies in melanoma. Vemurafenib was the first FDA-approved B-Raf^V600E^ mutant-specific inhibitor to show improved tumor reduction and progression-free survival compared to general chemotherapies (Chapman et al., 2011). A second FDA-approved B-Raf^V600E^ inhibitor Dabrafenib produced fewer side effects and had higher potency compared to Vemurafenib (Menzies et al., 2012). However, a significant challenge with these B-Raf^V600E^ targeted therapies is the acquisition of drug resistance by the tumor cells and reactivation of the MAPK pathway within the typical treatment course of 12 months (Czarnecka et al., 2020).

Given prolonged B-Raf^V600E^ inhibition resulted in reactivation of the MAPK pathway, MEK inhibitors such as Trametinib (Eroglu and Ribas, 2016) and Combimetinib (Ascierto et al., 2016) were developed to reduce drug resistance and increase treatment efficacy compared to B-Raf inhibitors alone. The MEK inhibitors used in combination with B-Raf inhibitors showed five-year progression-free survival of melanoma patients at 49% among patients with complete response and 16% among patients with partial response (Robert et al., 2019). Despite significant reduction in tumor burden, most patients relapsed with resistant melanoma tumors. Although immunotherapies such as anti-PD1 and anti-CTLA-4 (Hodi et al., 2010; Larkin et al., 2015) increasingly offer alternative therapeutic approaches, but are not universally effective, and resistance remains a persistent challenge (Sharma et al., 2017).

Common mechanisms leading to MAPKi-resistance involve MAPK reactivation through B-Raf^V600E^ copy-number amplification (Shi et al., 2012), reactivation of ERK (Corcoran et al., 2010), mutations in RAS genes (Nazarian et al., 2010; Romano et al., 2013) or suppression of neurofimbrin 1 (NF1) (Whittaker et al., 2013). These alterations lead to reactivation of MAPK and PI3K signaling pathways. Other resistance mechanisms involve subtype switching between the AXL and MITF subtypes (Johannessen et al., 2013; Konieczkowski et al., 2014; Müller et al., 2014; Smith et al., 2016) and epigenetic reprogramming (Emert et al., 2021; Jin et al., 2015; Khaliq and Fallahi-Sichani, 2019; Shaffer et al., 2017).

Recent evidence underscores the significance of downstream signaling changes, which occur in response to genetic mutations and transcriptional reprogramming and their potential dependence on alterations in cell cytoskeleton. For instance, in MAPKi-treated melanoma cells with hyperactivating Rac1^P29S^ in addition to B-Raf^V600E^ triggers an acute cell morphological response characterized by the development of extended lamellipodia containing dense actin networks. These structures serve to sequester and inactivate the tumor suppressor Merlin, thereby sustaining proliferation (Mohan et al., 2019). Furthermore, in B-Raf mutant melanoma cells that develop resistance after prolonged exposure to MAPKi display F-actin remodeling within the context of YAP/TAZ-dependent B-Raf inhibitor resistance (Kim et al., 2016). Misek *et al*. later revealed that these F-actin structures are reliant on shifts in RhoA signaling downstream of transcriptional effectors such MRTF and YAP1 (Misek et al., 2020). Finally, Orgaz *et al*. reported cytoskeletal remodeling and changes in expression and activity of the ROCK-myosin II pathway are signature events during the development of MAPKi resistance. Notably, ROCK-myosin II inhibition specifically kills resistant cells via intrinsic lethal reactive oxygen species (ROS) after a 72h exposure to ROCK inhibitors (Orgaz et al., 2020), likely in association with alteration in metabolic pathways (Boveris and Chance, 1973; Mullarky and Cantley, 2015; Nègre-Salvayre et al., 1997). However, the precise link between RhoA–ROCK activation and the induction of resistance-associated metabolic adaptations remain to be established.

Several studies have reported a direct functional interaction between remodeling of the cytoskeleton and metabolic shifts (Hu et al., 2016; Luther and Lee, 1986; Real-Hohn et al., 2010; Roberts and Somero, 1987; Silva et al., 2004). This suggests a plausible model for RhoA’s involvement in metabolic reprogramming through its role as a promoter of actomyosin structure assembly and activation. Specifically, F-actin bundles bind and stabilize the active form of the rate-limiting glycolytic enzyme phosphofructokinase (PFK)-1 (Luther and Lee, 1986; Roberts and Somero, 1987), while also moderating PFK degradation through sequestration of the E3-ubiquitin ligase Trim-21 (Park et al., 2020). Moreover, the glycolytic enzyme aldolase also binds to F-actin, and its release of aldolase from actin can be triggered by insulin (Hu et al., 2016). Building on these observations, our hypothesis posits that the resensitization of MAPKi-resistant melanoma to ROCK inhibition may be linked to the destabilization of F-actin structures followed by dampening metabolic activity, particularly glycolysis.

In this study we describe the generation of F-actin bundles through elevated RhoA activation in MAPKi-resistant melanoma cells. Contrary to expectation, however, enhanced glycolysis in these cells was independent of F-actin bundles. Instead, we found that RhoA activation altered metabolism through elevated glucose uptake because of elevated expression of the glucose transporter, GLUT1. Targeting the MAPKi-resistant cells with ROCK inhibitors reduced the glucose uptake and metabolism. We thus provide a new link between a well-described adaptation in Rho GTPase signaling in MAPKi drug resistant melanoma and metabolic adaptation.

## Results

### Prolonged exposure to MAPKi confers drug resistance in melanoma cells

To test the interplay between RhoA-ROCK signaling, F-actin organization, and metabolism in MAPKi-resistant melanoma, we generated five drug resistant melanoma cell lines *in vitro* by prolonged exposure (∼ 2 months) to MAPKi combination treatment of 100nM Dabrafinib + 10nM Trametinib (Supp. Fig. 1A). Following an initial reduction in proliferation, the resistant populations eventually restored their proliferation to similar levels to those of the DMSO treated control (naïve) population (Fig. 1A, Supp. Fig. 1B).

**Figure 1.**
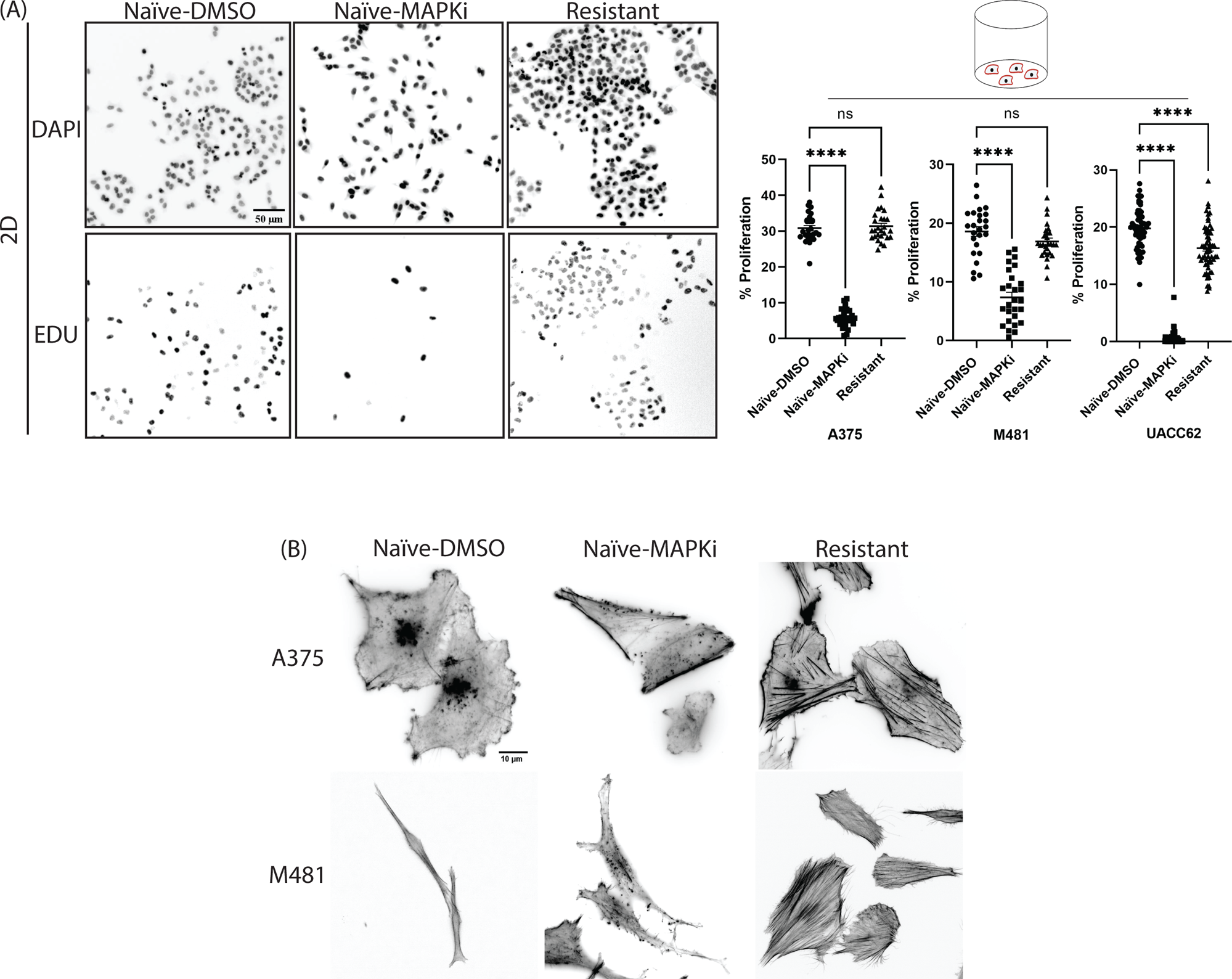
Melanoma cells acquire a proliferative phenotype upon prolonged exposure to MAPKi. (A) Flow chart to generate MAPKi-resistant melanoma cells. The naïve B-Raf V600E mutant melanoma cells upon exposure to MAPKi show an initial reduction in proliferation. But upon exposure to MAPKi for ∼2 months results in acquisition of resistance. These resistant cells were then cultured in the presence of MAPKi. (B) A375, M481 and UACC62 naïve and resistant cells were compared for proliferation using an imaging based EdU incorporation assay in 2D cell culture conditions. The naïve cells were treated with vehicle control or with MAPKi and the proliferation compared to MAPKi-resistant melanoma cells. The % Proliferation was determined as the ratio of EdU-positive cells to the total cells determined by DAPI counterstain. The scale bar on the image is 50µm. (C) A375 naïve and resistant cells were cultured on glass coverslips. The naive cells were either treated with vehicle control or MAPKi. All the cells were then fixed and stained with phalloidin to evaluate F-actin structures. The scale bar on the image is 10µm. For (B) Three biological repeats were performed with a minimum of 9 images acquired per sample per repeat. The statistical analysis was performed using one-way ANOVA. For (C) Imaging was performed for three biological repeats with a minimum of 10 images acquired per sample per biological repeat. For the statistical significance; ns P ≥ 0.05, *P ≤ 0.05, **P ≤ 0.01, ***P ≤ 0.001, ****P ≤ 0.0001. The data represented in (B) is mean ± s.e.m.

Prior work has underscored the importance of microenvironmental stiffness in determining therapeutic efficacy (Mohan et al., 2019; Murali et al., 2019). To validate that the MAPKi resistance observed in these melanoma cells was not contingent on the microenvironment, we performed proliferation assays also on soft collagen gels. Naïve cells exhibited similar reduction in proliferation upon acute treatment (48 hours) to MAPKi as on plastic substrates, whereas MAPKi-resistant melanoma cells retained their proliferation level (Supp. Fig. 1C). This confirmed the independence of acquired resistance to MAPKi from the microenvironment.

In prior studies, other groups had reported alterations in F-actin structures upon acquisition of resistance to B-Raf inhibitors such as PLX4720 (Kim et al., 2016; Misek et al., 2020). This led us to investigate whether our MAPKi combination treatment would induce analogous cytoskeletal response. Acute MAPKi treatment resulted in mild but prominent peripheral F-actin bundle formation. This response was greatly amplified upon resistance acquisition. The resistant cell populations displayed thick F-actin fibers cross-cross the entire cytoplasm (Fig. 1B, Supp. Fig. 1D), aligning with the notion that MAPKi resistance may relate to shifts in F-actin organization.

### Drug resistant melanoma cells acquire an actin-independent metabolic state

Next, we determined the glycolytic (ECAR) and mitochondrial respiration (OCR) rates to test whether resumption of proliferation in MAPKi-resistant cells correlated with shifts in metabolic activity when compared to both naïve vehicle control and those subjected to acute MAPKi-treatment. The naïve cells exposed to acute MAPKi treatment showed a strong reduction in ECAR, though OCR remained unaffected (Fig. 2A). In contrast, the MAPKi-resistant melanoma cells not only restored ECAR levels similar to a state similar to that of naïve cells under vehicle control but also exhibited a significant increase in OCR. This metabolic shift might be associated with changes in focal adhesion (FA) organization linked to the bundling of F-actin (Ciobanasu et al., 2012; Kim et al., 2012; Romani et al., 2021), which has been shown to tether mitochondria and promote their fusion (Daniel et al., 2019). Based on this we evaluated FA in naïve and MAPKi-resistant cells. Immunofluorescence staining for phospho-tyrosine (pY) and phalloidin showed the MAPKi-resistant melanoma cells had larger adhesions, in line with the increased density of F-actin fibers (Supp. Fig 2A).

**Figure 2.**
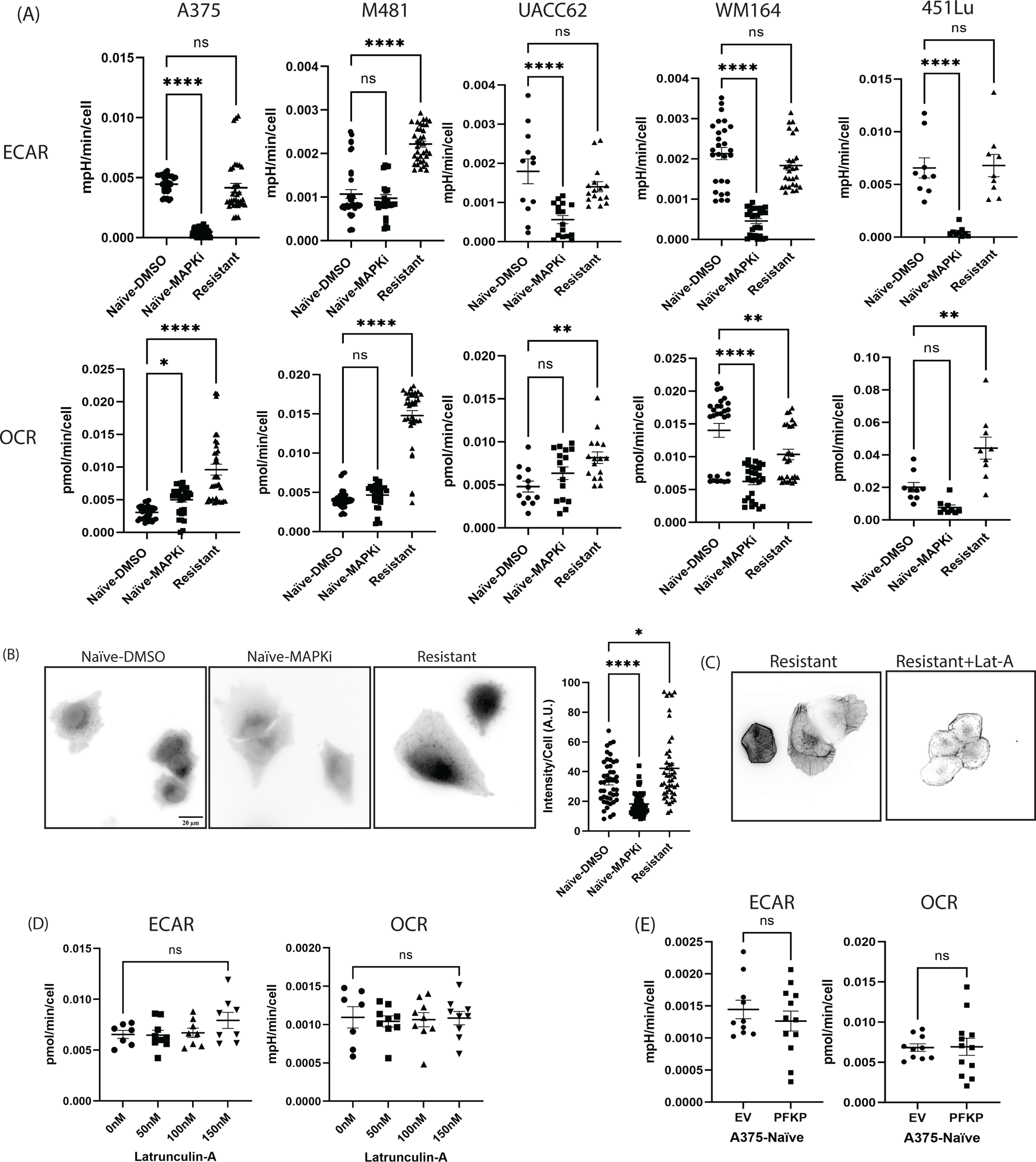
Metabolic characterization of resistant melanoma cells. (A) A375, M481, UACC62, WM164 and 451Lu were evaluated for the metabolic state. The resistant melanoma cells were compared to naïve cells with vehicle control or MAPKi treatment and seahorse assay performed to evaluate ECAR (readout of glycolysis) and OCR (readout for oxidative phosphorylation). The metabolic rates, ECAR and OCR were normalized to per cell through fixation, DAPI staining and cell count using ImageJ after seahorse assay. (B) Glucose uptake in naïve and resistant A375 melanoma cells was evaluated using NBD-glucose. The cells were then imaged, and the image intensity was quantified per cell using ImageJ. (C) To evaluate the role of F-actin structures in metabolism the A375 resistant melanoma cells were treated with Latrunculin-A to evaluate actin depolymerization. (D) Seahorse assay was then performed for the resistant cells treated with Latrunculin-A after 4h of treatment. (E) A375 naïve cells were overexpressed with a PFKP-GFP followed by seahorse assay. All experiments were performed with 3 biological repeats. For (A) A375, M481, and WM164 had 9 data points per repeat. The UACC62 and 451Lu had atleast 3 datapoint per repeat. For (B) Each biological sample in the repeat had at least 12 images. (D) and (E) had 3 data points per repeat. The statistical analysis was performed using one-way ANOVA for (A), (B) and (D). Welch’s t-test was used for (E). For the statistical significance; ns P ≥ 0.05, *P ≤ 0.05, **P ≤ 0.01, ***P ≤ 0.001, ****P ≤ 0.0001. The data represented in (B) is mean ± s.e.m.

To corroborate the observed increased metabolic activity, we performed unbiased metabolomics for the naïve cells treated with vehicle control or acute MAPKi and on resistant melanoma cells (Supp Table 1 and Supp Fig 2C). The results of glycolysis and TCA cycle confirmed a resumption in glucose uptake, glycolysis, and TCA cycle in MAPKi-resistant melanoma cells (Supp Fig 2D). To validate the prediction from unbiased metabolomics of elevated glucose uptake in MAPKi-resistant cells, we directly measured glucose uptake rates using the fluorescent glucose analog NBD-glucose. While naïve cells exhibited a significant reduction in glucose uptake upon acute MAPKi treatment as previously reported (Parmenter et al., 2014) (Fig 2B). In contrast, MAPKi-resistant melanoma cells maintain ed or even slightly increased the glucose uptake relative to vehicle control treated naïve cells (Fig 2B) potentially explaining why these cells can resume ECAR.

A parallel mechanism contributing to the sustained glucose uptake in MAPKi-resistant melanoma cells may relate directly to the significant increase in F-actin bundling (Fig. 1C). Park *et al*. (Park et al., 2020) reported that in human bronchial epithelial cells (HBECs) and lung cancer cells cortical F-actin bundles can sequester the E3 ubiquitin ligase TRIM21, thereby reducing the degradation of the rate-limiting glycolytic enzyme phosphofructokinase (PFK). In that study, depolymerization of F-actin structures was sufficient to restore TRIM21’s function in polyubiquitinating PFK, followed by significant reduction in glycolytic activity. Thus, we hypothesized that F-actin depolymerization in MAPKi-resistant melanoma cells would reduce ECAR, and potentially re-sensitize cells to MAPKi treatment. Accordingly, we treated MAPKi-resistant melanoma cells with Latrunculin-A (Lat-A), a potent F-actin depolymerizer, and found by imaging a marked reduction in F-actin structures (Fig 2C). Contrary to our expectation, both ECAR and OCR were unaffected by the disruption of F-actin bundles (Fig 2D). We further tested whether overexpression of PFK in naïve melanoma cells would increase glycolysis and thus potentially induce MAPKi-resistance. In line with the negative result of F-actin abrogation elevated PFK had no effect on metabolism (Fig 2E). Hence, the glycolytic shifts observed in MAPKi-resistant melanoma cells are unlikely to relate to the TRIM-21/PFK mediated metabolic shifts found in HBECs. Although increased F-actin bundling, and contraction are morphological signatures of MAPKi-resistant cells they appear not to be causal for the maintenance of glucose metabolism associated with MAPKi resistance.

### RhoA activation in drug resistant melanoma cells promotes elevated glucose uptake and metabolism

Misek *et al*. reported an activation of RhoA signaling in B-Raf mutant melanoma cells (Misek et al., 2020) and Orgaz *et al*. showed that treatment of MAPKi-resistant melanoma cells with Rho-activated kinase inhibitors such as GSK269962A along with immunotherapies significantly reduced tumor burden (Orgaz et al., 2020). Both observations conceptually align with our observation of increased F-actin bundling in MAPKi-resistant cells. Therefore, we wondered whether our panel of resistant cells may also be subject to elevated RhoA activity, possibly linked to metabolism. We performed GTP-RhoA pull downs using Rhotekin beads and found an increased activation of RhoA in MAPKi-resistant melanoma cells compared to their naïve counterparts (Fig 3A). This increase in RhoA activity was accompanied by an increased expression of total RhoA in resistant cells (Fig 3A). Accordingly, upon normalizing the active RhoA to total RhoA the relative RhoA activity was no longer significantly increased in MAPKi-resistant cells (Supp. Fig 3A). Hence, the higher levels of active RhoA relate primarily to an increased expression of RhoA protein in MAPKi-resistant melanoma cells. To further validate our result, we examined RhoA activation by measuring the phosphorylation state of myosin light chain 2 (MLC2). In support of elevated RhoA signaling in MAPKi-resistant cells phosphorylated MLC2 (pMLC2) was higher in MAPKi-resistant cells compared to naïve (Supp Fig. 3B). This was accompanied by a modest increase in total MLC2. However, normalization of pMLC2 to MLC confirmed a more than two-fold increase in MLC2 phosphorylation, which underlines the robust elevation of RhoA signaling in MAPKi-resistant cells.

**Figure 3.**
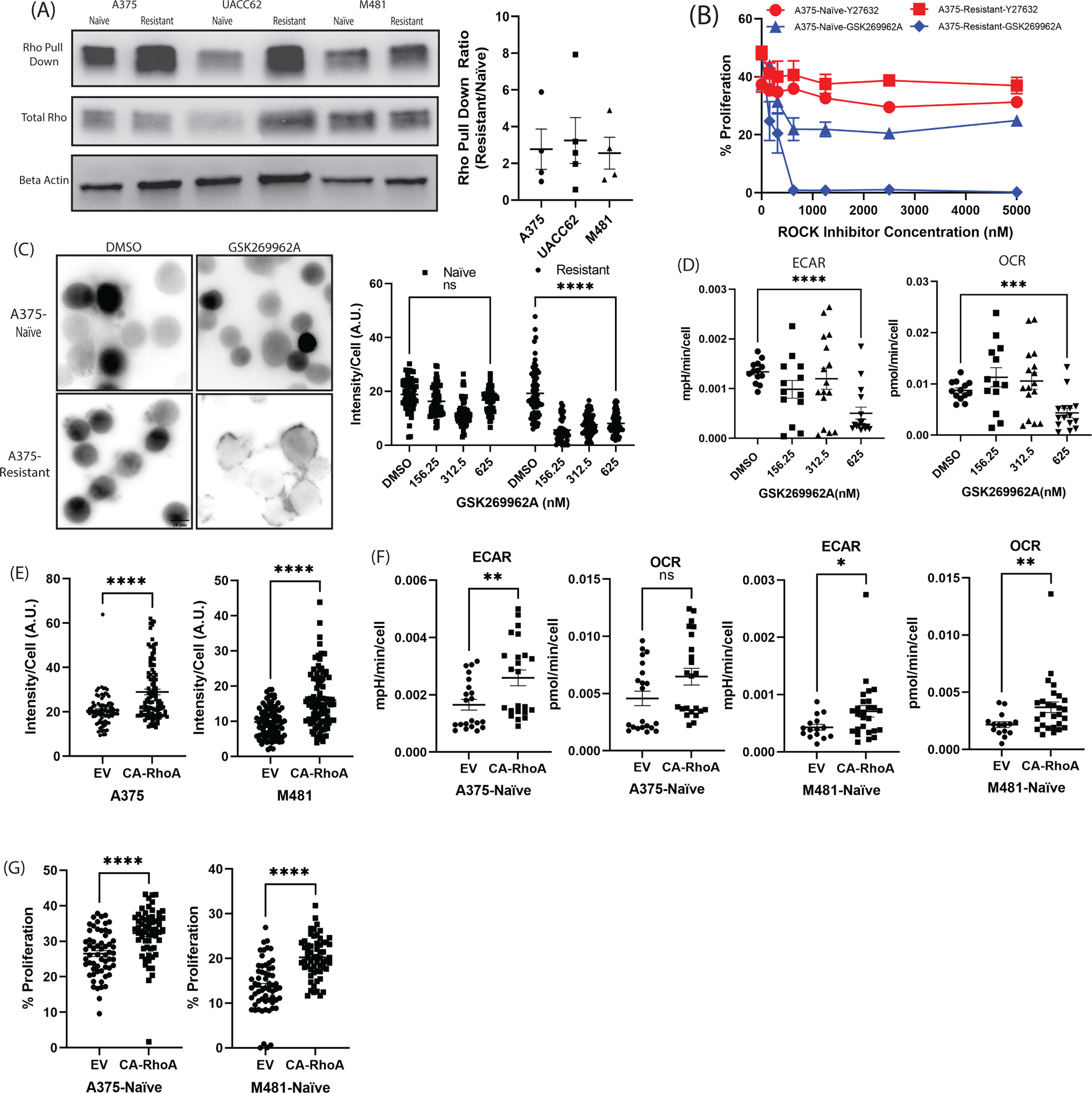
Targeting RhoA targets metabolism in MAPKi-resistant melanoma cells. (A) RhoA activation was evaluated in A375, UACC62 and M481 naïve and resistant melanoma cells using the RhoA activation pull down assay. The band intensities were quantified for the rho pull down as a ratio of resistant to naïve cells. (B) The A375 naïve and resistant cells were treated with Y27632 and GSK269962A for 48h followed by EdU incorporation assay to evaluate proliferation. (C) A375 naïve and resistant melanoma cells treated with GSK269962A were evaluated for glucose uptake using NBD-glucose and the intensities quantified per cells using ImageJ. (D) Metabolic rate was quantified for A375 resistant melanoma cells treated with GSK269962A and evaluated for ECAR and OCR. (E) RhoA was activated in naïve A375 and M481 using a constitutively active RhoA (CA-RhoA) and evaluated for glucose uptake using NBD-glucose assay and image intensities quantified per cell which was compared to empty vector (EV). (F) Seahorse assay was performed to evaluate ECAR and OCR in naïve A375 and M481 melanoma cells with EV or CA-RhoA. (G) Proliferation between EV and CA-RhoA was evaluated using EdU incorporation assay for the naïve A375 and M481 melanoma cells. For (A) 4 biological repeats, (B) Three biological repeats were performed with three data points per repeat, (C) (E) and (G) Three biological repeats with at least 10 images per sample per repeat, (D) Three biological repeats with four data points per sample per repeat. (F) Three biological repeats with six data points per repeat. The statistical analysis was performed using one-way ANOVA for (C), and (D). Welch’s t-test was used for (E), (F) and (G). For the statistical significance; ns P ≥ 0.05, *P ≤ 0.05, **P ≤ 0.01, ***P ≤ 0.001, ****P ≤ 0.0001. The data represented in (B) is mean ± s.e.m.

If RhoA activation indeed promotes MAPKi resistance via RhoA-ROCK signaling, as suggested by the experiments reported by Orgaz *et al*. (Orgaz et al., 2020) and Misek *et al*. (Misek et al., 2020), then ROCK inhibitors will specifically affect the MAPKi-resistant cell population and less the naïve melanoma cells. We used the ROCK inhibitors Y27632 and GSK269962A in the naïve and MAPKi-resistant melanoma cells. While Y27632-treatment had no effect, application of GSK269962A (ROCKi) showed a robust dose response in proliferation below 625nM concentration in MAPKi-resistant cells (Fig 3B, Supp. Fig. 3C). At 625nM ROCKi proliferation was completely blocked in these cells. Importantly, naïve cells exhibited a partial response, but proliferation plateaued at concentrations greater than 625nM (Fig 3B), indicating that proliferation in MAPKi-resistant melanoma cells is specifically driven by ROCK activity. This was corroborated by analysis of cytotoxic effects of ROCKi using Ethidium homodimer-based imaging. Upon treatment with ROCKi, MAPKi-resistant melanoma cell death was less than 10% (Supp Fig. 3D).

Next, we examined whether the effect of ROCKi treatment on MAPKi-resistant melanoma cell proliferation would relate to metabolic activity. We first evaluated the effect on glucose uptake. We treated both the naïve and MAPKi-resistant melanoma cells with vehicle (DMSO) control and ROCKi. There was no significant change in glucose uptake by the naïve cells (Fig. 3C). The MAPKi-resistant cells, however, showed a strong reduction in glucose uptake upon application of ROCKi (Fig. 3C). Second, ROCKi treatment of MAPKi-resistant cells also decreased ECAR and OCR (Fig. 3D, Supp. Fig 3E). Together, these experiments provided first indication of a possible link between RhoA activation and metabolism in MAPKi resistant cells. The link can be intercepted by ROCKi.

To test the role of RhoA activation in glucose uptake and metabolism more directly and to rule out the possibility that ROCKi’s effect on glucose uptake in resistant cells is merely the result of dampened cell growth, we expressed constitutively active RhoA (CA-RhoA) in naïve melanoma cells. We validated the overall activation of RhoA in these cells by the increase in phosphorylated MLC (p-MLC) (Supp Fig 3F). The gain-of-function in RhoA induced increased glucose uptake (Fig. 3E), elevated ECAR and OCR values (Fig. 3F & Supp. Fig. 3G) and increased proliferation (Fig 3G & Supp. Fig. 3H). However, treatment of these CA-RhoA cells with MAPKi did not yield resistance (Supp. Fig. 3I). Khosravi-Far *et al*. had previously reported that the activation of RhoA through lab engineered mutations such as G14V or Q63L may not be enough to induce such transformations by themselves (Khosravi-Far et al., 1995) and that activation of RhoA alone is not sufficient to induce transformation of naïve cells. However, they play a crucial role in promoting cell cycle progression. Overall, these results indicate that activation of RhoA is sufficient to increase glucose uptake and elevate metabolism and thus to promote cell proliferation.

### PKA activation results in RhoA inactivation to reduce metabolism and proliferation

Activation and inhibition of RhoA is mediated through the tightly regulated cycling of accessory proteins: guanine nucleotide dissociation inhibitor (GDI), which retains Rho in a GDP bound (inactive) state; GTPase activating proteins (GAPs), which stimulate hydrolysis of GTP to GDP; and guanine nucleotide exchange factors (GEFs), which activate GTPase to release the GDP bound to RhoA (Ridley and Hall, 1992). One of the factors that mediates RhoA activation/inactivation is cyclic AMP-dependent protein kinase A (PKA) (DerMardirossian and Bokoch, 2005; Qiao et al., 2003; Tkachenko et al., 2011). PKA plays a role in diverse cellular activities and is reported to exhibit negative temporal and spatial correlation with the activation of RhoA (Tkachenko et al., 2011). The correlation depends on the phosphorylation of RhoA (Ser 188) by PKA, an event known to facilitate binding of RhoA to Rho GDI leading to RhoA displacement and inhibition of RhoA interaction with downstream effectors (Lang et al., 1996). We hypothesized that an increased activation of RhoA in MAPKi-resistant melanoma cells could be an effect of reduced activation of PKA. Indeed, comparison of PKA activation in naïve and MAPKi-resistant melanoma cells using a colorimetric assay indicated reduced activation of PKA in MAPKi-resistant melanoma cells (Fig 4A).

**Figure 4.**
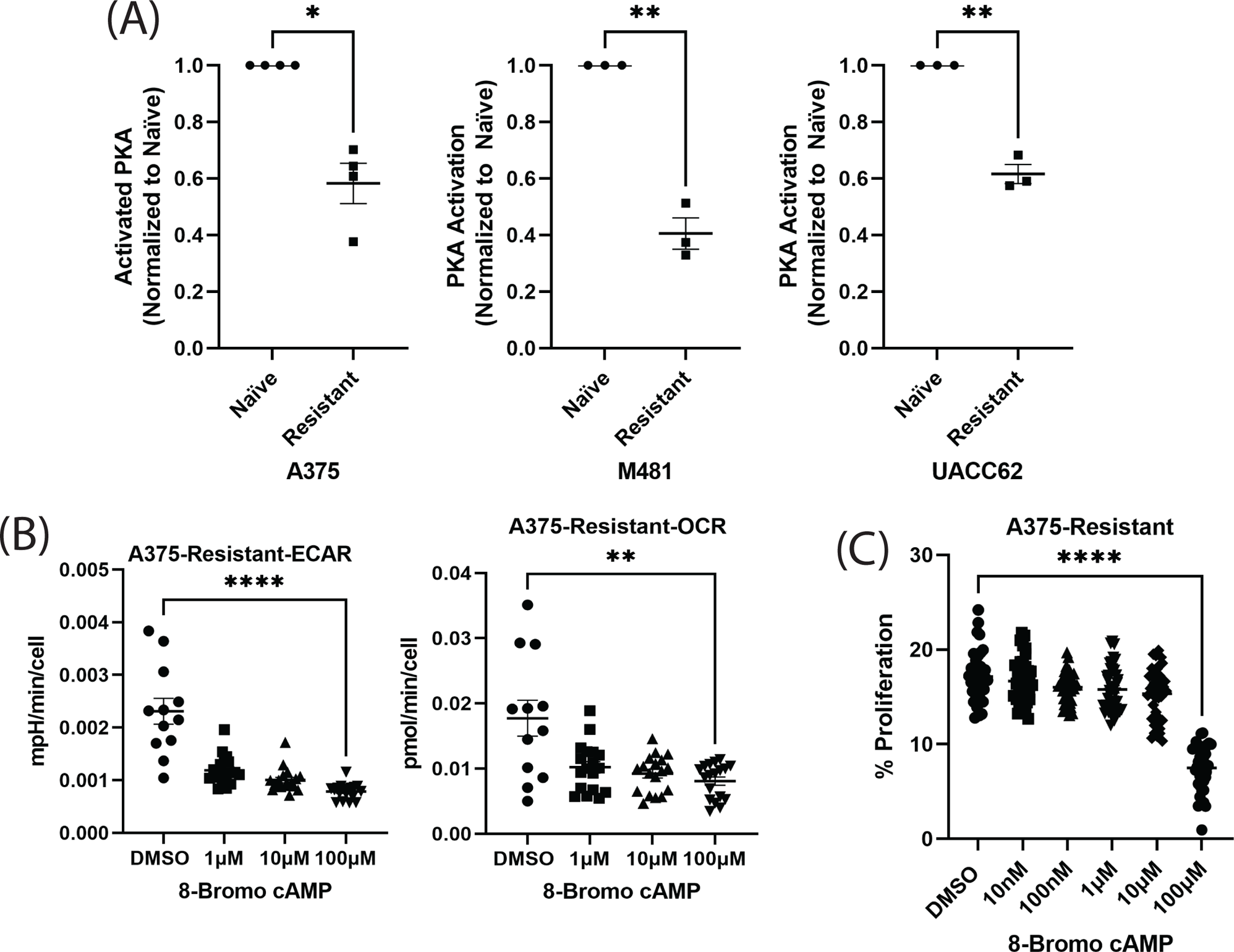
PKA activation in resistant melanoma cells alters metabolism and proliferation. (A) The activity of PKA in A375, M481 and UACC62 naïve and resistant melanoma cells was evaluated using the PKA activation assay. (B) To induce PKA activation in A375 resistant cells 8-Bromo-cAMP was used and the metabolic state was evaluated using seahorse assay. (C) Proliferation was evaluated in A375 resistant melanoma cells upon 8-Bromo-cAMP treatment using EdU incorporation assay. For (A) each data point represents a biological repeat. For (B) Three biological repeats were performed with four data points per repeat. For (C) three biological repeats ten data points per repeat. The statistical analysis was performed using one-way ANOVA for (B) and (C). Welch’s t-test was used for (A). For the statistical significance; ns P ≥ 0.05, *P ≤ 0.05, **P ≤ 0.01, ***P ≤ 0.001, ****P ≤ 0.0001. The data represented in (B) is mean ± s.e.m.

Based on this result, we expected that inducing PKA activation should reduce ECAR and OCR in MAPKi-resistant cells. To evaluate this, we treated MAPKi-resistant cells with 8-bromo-cAMP, a cell-permeable analog of cAMP that activates cyclin-AMP dependent protein kinase. As predicted, after 48 hours of treatment MAPKi-resistant cells showed a reduction in both ECAR and OCR in an 8-bromo-cAMP concentration-dependent manner (Fig 4B). The proliferation of the MAPKi-resistant cells treated with 8-bromo-cAMP was also reduced (Fig. 4C). These results further supported our model that elevated RhoA activation in MAPKi-resistant cells promotes a metabolic shift in support of sustained proliferation.

### RhoA activation promotes glucose uptake through an IRS1-P85 pathway

How is elevated RhoA activation linked to glucose uptake and metabolism? The interactions of glucose and insulin, specifically binding to insulin receptor, leads to phospho-activation of insulin receptor substrate (IRS) proteins. Phosphorylated IRS proteins act as a docking site for the p85 regulatory subunit of PI3K, which in turn leads to PI3K activation to promote glucose uptake in skeletal muscle and adipose tissue by glucose transporter 4 (GLUT4). Noboru *et al*. reported that in adipose cells ROCK regulates insulin-stimulated glucose transport via PI3K activation downstream of IRS1 phosphorylation (Furukawa et al., 2005). Based on these observations, we hypothesized that MAPKi-resistant cells stimulate IRS1 through the activation of RhoA-ROCK. Hence, targeting MAPKi-resistant cells with the ROCKi would result in reduced phosphorylation of IRS1 (pIRS1), concomitantly leading to reduced phospho-activation of P85 (pP85) and reduced glucose uptake.

To test this hypothesis, we first checked the baseline difference in expression of IRS1, pIRS1, P85, pP85, GLUT1, GLUT3 and GLUT4 between naïve and MAPKi-resistant melanoma lines. We observed increased expression of IRS1 in the MAPKi-resistant cells compared to the naïve cells. pIRS1 levels were similar between the two cell types. However, MAPKi-resistant cells had higher pP85 levels (Fig. 5A). Prior work elucidated the role of glucose transport through GLUT4 upon activation of RhoA (Furukawa et al., 2005). In our experiments, western blot analysis for total protein expression showed no statistical difference in protein expression for GLUT1, GLUT3 and GLUT4 between the naïve and MAPKi-resistant cells (Fig. 5A). However, GLUT1 and GLUT4 did show a trend towards increase in total protein expression in the resistant cell population. Koch *et al*. reported an enhanced expression of GLUT1 linked to melanoma malignancy (Koch et al., 2015). Thus, we sought to corroborate the trend of increased GLUT1 and GLUT4 expression by flow cytometry measuring GLUT expression on the cell surface. Indeed, this confirmed higher GLUT1 and GLUT4 surface expression in resistant cells (Fig. 5B and Supp. Fig. 4), along with GLUT3. Hence, MAPKi-resistant cells seem to amplify proliferation-promoting glucose uptake by higher surface density of GLUT.

**Figure 5.**
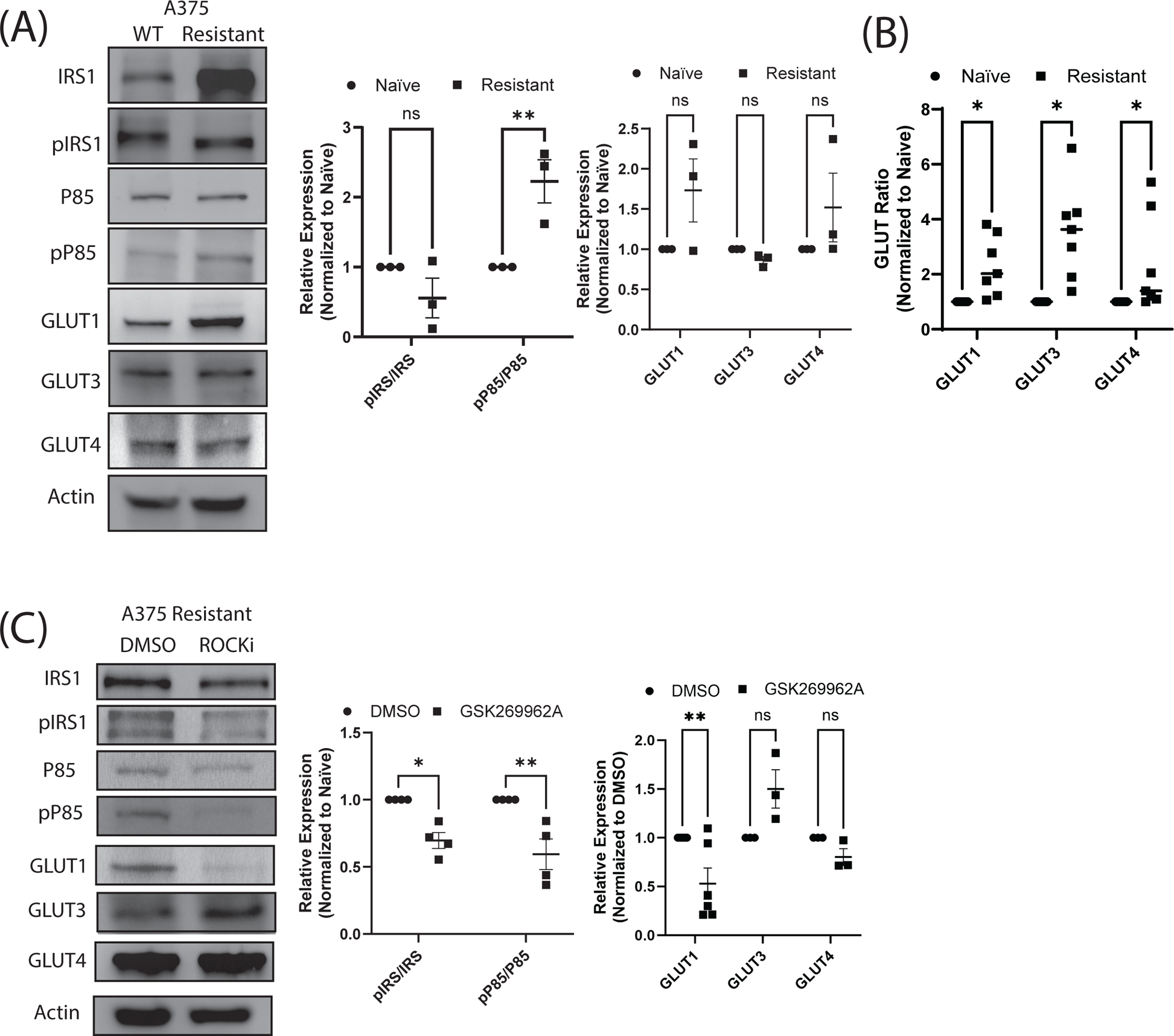
Resistant melanoma cells phosphor activate IRS1, p85 to elevate glucose uptake through GLUT1. (A) Western blots were performed to compare the A375 naïve and resistant melanoma cells for the expression of IRS1, p85, GLUT1 GLUT3 and GLUT4. (B) Flow cytometry results to compare GLUT1, GLUT3 and GLUT4 between naïve and resistant melanoma cells. (C) Protein expression change for IRS1, p85 and GLUT1 GLUT3 and GLUT4 were evaluated upon ROCKi GSK269962A treatment using western blots. (D) Flow cytometry to evaluate change in GLUT1, GLUT3 and GLUT4 for MAPKi resistant melanoma cells upon ROCKi treatment compared to vehicle control. For both (A) and (C) the expression of the phospho proteins of IRS1 and p85 were ratioed to the total IRS1 and p85. All proteins were ratioed to their actin loading controls. For (A), (B), (C) and (D) each data point represents a biological repeat. The statistical analysis was performed using Two-way ANOVA. For the statistical significance; ns P ≥ 0.05, *P ≤ 0.05, **P ≤ 0.01, ***P ≤ 0.001, ****P ≤ 0.0001. The data represented in (B) is mean ± s.e.m.

To test whether the GLUT expression increase in melanoma relates to the pathway Furukawa et al. described in adipocytes, we evaluated GLUT protein expression under ROCKi. In this case we found a strong and selective reduction of GLUT1 expression. Since GLUT1 is known to be expressed only at the cell surface the western blot analysis indicates that ROCKi reduces the capacity of glucose uptake primarily by lowering the glucose uptake through GLUT1 (Fig. 5C). In summary, these results demonstrate that elevated metabolism in MAPKi-resistant melanoma cells is a consequence of ROCK activation, which promotes GLUT1 expression and glucose uptake via a signaling cascade in which translates Rho kinase activity into PI3 kinase activity.

## Discussion

Numerous studies have identified shifts in RhoA expression and activation as hallmark features of increasingly aggressive cancers (Crosas-Molist et al., 2022). The heightened RhoA signaling has been linked to the acquisition of an amoeboid cell phenotype, which is particularly effective in invasion, survival, and metastatic spread (Crosas-Molist et al., 2022; Farrugia et al., 2020; Orgaz et al., 2020). Amoeboid-based migration, which can be either protease-dependent (Orgaz et al., 2014) or independent (Driscoll et al., 2022; Sahai and Marshall, 2003; Wolf et al., 2003), is characterized by high actomyosin contractility, with blebs being the prominent functional protrusions (Sahai and Marshall, 2003). Recent studies by Remi *et al*. show ROCK-Myosin II activity in pancreatic ductal adenocarcinoma (PDAC) supports transcriptional program conferring amoeboid invasive abilities (Samain et al., 2023). A point not discussed but shown in the paper through gene set enrichment analysis is the elevated glycolytic state of amoeboid cells.

Our study further identifies drug resistance as a distinctive characteristic of heightened RhoA signaling, in line with findings from Orgaz and Misek *et al*. (Misek et al., 2020; Orgaz et al., 2020). Melanoma cells resistant to MAPKi exhibited sustained proliferation when compared to the naïve acute MAPKi-treated melanoma cells. Previous research has indicated that ROCK and Myosin II regulate G2/M cell cycle progression and cytokinesis in MAPKi-resistant melanoma cells (Orgaz et al., 2020). However, a lingering question pertained as to how RhoA activation promotes MAPKi resistance in melanoma cells and the mechanism of action upon targeting these MAPKi-resistant cells with ROCKi.

We discovered that this elevated RhoA activation in MAPKi-resistant melanoma cells results in the activation of ROCK, which subsequently results in the pIRS1 activation to promote glucose uptake. Targeting ROCK with ROCKi resulted in diminished glucose uptake, metabolic activity, and proliferation. This phenomenon mirrors findings by Furukawa *et al*., who previously demonstrated how RhoA activation led to IRS1 Ser632/635 phosphorylation by ROCK in adipose cells to promote insulin dependent glucose uptake through GLUT4. In contrast, we propose that upregulated glycolysis in MAPKi resistant melanoma cells originates primarily in GLUT1, whose expression we see raised in both western blots and flow cytometry. Consistently, we see a reduction of GLUT1 expression upon ROCKi in MAPKi-resistant melanoma cells. However, these cells showed no change in GLUT3 or GLUT4 expression. This deviates from the work of Furukawa *et al*. where the glucose transport was attributed to shifts in GLUT4, however without presenting evidence that GLUT1 and GLUT3 are not affected (Furukawa et al., 2005). Moreover, their work did not evaluate the change in expression or surface localization of GLUT4. Thus, it is plausible that the strong shifts specifically in GLUT1’ we observed downstream of elevated RhoA/ROCK signaling in MAPKi-resistant cells would to some extent also be found in adipocytes. Regardless of apparent difference in the primary ROCK-targeted transporter isoform between adipocyte and melanoma, our data provide yet another example of cancer cells repurposing of cellular pathways from completely different contexts as a mechanism for acute self-defense against therapeutic insults.

In conclusion, our findings concerning the role of RhoA/ROCK in glucose uptake and metabolism underscore the potential clinical advantages of targeting melanomas through the RhoA/ROCK pathway. They also provide further evidence of the extensive roles that Rho family GTPases play in cancer cell processes, extending beyond their conventional function as regulators of the cytoskeleton. Given the multitude of regulators for Rho GTPases, it is plausible that a subset of specialized GEFs or GAPs modulate non-canonical functions, such as promoting glucose uptake, in specific contexts. These dedicated regulatory circuits may become promising candidates for RhoA-centric therapeutic strategies.

## Acknowledgements and Funding Sources

We thank Dana Kim Reed for technical support; Kushal Bhatt for assistance with plasmid design; Gabriel Muhire Gihana for assistance in optimizing myosin detection; Prapti Modi for providing editorial corrections on the manuscript, and Lauren Zacharias and the Children’s Research Institute’s metabolomic facility for metabolomics analysis. Ralph J. DeBerardinis is supported by the Howard Hughes Medical Institute and N.C.I. Grants R35CA22044901 and 2P50CA070907-21A1. Work in the Danuser lab has been supported by the National Institute of General Medical Sciences grant R35 GM136428 and by the Welch Foundation grant I-1840.

---

**Supplemental Figure 1. Resistant melanoma cells proliferation in microenvironment independent.**

(A) WM164 and 451Lu naïve and resistant cells were compared for proliferation using an imaging based EdU incorporation assay in 2D cell culture conditions. The naïve cells were treated with vehicle control or with MAPKi and the proliferation compared to MAPKi-resistant melanoma cells. The % Proliferation was determined as the ratio of EdU-positive cells to the total cells determined by DAPI counterstain. (B) A375, M481 and WM164 naïve and resistant cells were compared for proliferation using an imaging based EdU incorporation assay with the cells cultured on bovine soft collagen. The naïve cells were treated with vehicle control or with MAPKi and the proliferation compared to MAPKi-resistant melanoma cells. Maximum intensity Z-projections were generated for cells cultured on collagen and proliferation evaluated. The % Proliferation was determined as the ratio of EdU-positive cells to the total cells determined by DAPI counterstain. For (A) Three biological repeats were performed with a minimum of 9 images acquired per sample per repeat. (B) Three biological repeats were performed with a minimum of 6 images acquired per sample per repeat. The statistical analysis was performed using one-way ANOVA. For the statistical significance; ns P ≥ 0.05, *P ≤ 0.05, **P ≤ 0.01, ***P ≤ 0.001, ****P ≤ 0.0001. The data represented in (B) is mean ± s.e.m.

**Supplemental Figure 2. Morphological and metabolic characterization of naïve and resistant melanoma cells.**

(A) A375 naïve and resistant cells were cultured on glass cover slips to evaluate the difference in focal adhesions and colocalization with F-actin structures. The cells were fixed with immunofluorescence performed for phosphor-tyrosine (pY) and counterstained with phalloidin. (B) Vinculin expression was also evaluated using western blots for the A375 naïve cells treated with vehicle or MAPKi and compared to resistant cells. (C) To validate seahorse results, unbiased metabolomics was performed for A375 naïve cells treated with vehicle or MAPKi were compared to resistant cells. The heatmap results show the complete panel of metabolomics results. (D) Specific metabolite results from unbiased metabolomics for glycolysis and TCA-cycle. (E) Principal component analysis for A375 naïve cells treated with vehicle or MAPKi and resistant cells. For (A) and (B) Two biological repeats were performed with (A) having 10 images per sample per repeat. (C), (D) and (E) Three biological samples were generated but all the samples were run at the same time for unbiased metabolomics.

**Supplemental Figure 4. Characterizing glucose transporters in melanoma cells using flow cytometry.**

The results show the gating of negative cells. Negative gates were applied to cells with the staining for GLUT1 stained with Alexa Fluor 488, GLUT 3 with Alexa Fluor 568 and GLUT4 with Alexa Fluor 647.

## Materials and Methods

### Cell Lines

All melanoma cells described in this work harbor the B-Raf V600E mutation. A375 cells (human malignant melanoma cells harvested from a 54-year-old female CRL-1619, ATCC, RRID: CVCL_0132), UACC62 cells (human malignant melanoma ABC-TC534S, Accegen, RRID: CVCL_1780), M481 primary cells were acquired from the University of Michigan via Dr. Sean Morrison at UT Southwestern Medical Center, WM164 (metastatic human melanoma cell line established from a metastatic site in a 22-year-old male with stage IV superficial spreading melanoma WM164-01-0001, Rockland, RRID: CVCL_7928), 451Lu (xenograft aggressive metastatic sub-line of WM164; 451Lu-01-0001, Rockland, RRID: CVCL_6357).

### Tissue Culture Materials

All cells were cultured in incubators at 37°C with 5% CO_2_ and supplemented with 0.2% antibiotic-antimycotic purchased from Gibco (15240062). The A375 and UACC62 cells were cultured in Dulbecco’s modified eagle media (DMEM), with high glucose and L-glutamine, which was purchased from Gibco (11965-167) and supplemented with 10% fetal bovine serum (FBS). The M481 primary cells were cultured in Dermal Basal Medium purchased from ATCC (PCS-200-030) and supplemented with adult melanocyte growth kit purchased from ATCC (PCS-200-042) which includes rh-insulin, ascorbic acid, L-Glutamine, epinephrine, CaCl_2_, peptide growth factor and M8 supplement. The WM164 and 451Lu cells were cultured in MCDB-L15 which is a combination of MCDB153 containing L-glutamine and 28mM HEPES without sodium bicarbonate, purchased from Sigma Aldrich (M7403) and Leibovitz’s L-15 media which was purchased from Sigma Aldrich (L1518). The MCDB powder was re-suspended in 900mL double distilled water. 1.2g cell culture grade sodium bicarbonate and adjust pH to 7.6 with 5M NaOH using a pH meter. The volume was brought to 1L and filter sterilized with a 0.22µm filter. The combination media was made with 80% final concentration of MCDB153, 20% Leibovitz’s L-15, 1.68mM CaCl_2_, and 2% FBS and then filtered with a 0.22µm filter. Phenol red-free DMEM with high glucose and L-glutamine for fluorescence imaging was purchased from Thermo Fisher Scientific (21063-045). The phenol red free media was supplemented with 10% FBS. Trypsin/EDTA was purchased from Thermo Fisher Scientific (R001100). Trypsin Neutralizer (TN) was purchased from Thermo Fisher Scientific (R-002-100). For culturing cells on collagen, bovine collagen I with a molecular weight of ∼300kDa was purchased from Advanced Biomatrix (5005-100). Dimethyl Sulfoxide (DMSO) was purchased from Thermo Fisher Scientific (BP231-100). Dabrafenib (GSK2118436), a BRAF V600-targeting inhibitor (S2807), Trametinib (GSK1120212), a potent MEK1/2 inhibitor (S2673), Y27632 (S1049) and GSK269962A (S7687), selective ROCK1 and ROCK2 inhibitor were purchased from Selleckchem. Ethidium Homodimer (EtHd), a cell impermeant viability marker, was purchased from Thermo Fisher Scientific (E1169). Hoechst 33342, nuclear stain was purchased from molecular probes (H3570). Click-iT based imaging assay to evaluate proliferation was performed using ascorbic acid (A92902), copper sulfate (PH1477) which were purchased from Millipore Sigma and Alexa Fluor Azide (B6830) was purchased from Lumiprobe. To evaluate glucose uptake, 2-deoxy-2-[(7-nitro-2,1,3-benzoxadiazol-4-yl)amino]-D-glucose (NBD-Glucose) was purchased form Cayman chemical (11046).

### Generating drug resistant melanoma cells

Naïve melanoma cells unexposed to MAPK inhibitor combination of Dabrafenib and Trametinib were setup in 10cm dishes. Cells were grown to ∼70% confluency then washed with 1X PBS, trypsinized, and harvested by centrifugation at 300g for 5 mins. The pelleted cells were resuspended in 5mL media, and their concentration determined using cell counter (Nexelcom, Cellometer Auto 1000). Three 10-cm plates were setup for the naïve cells and seeded with 2 million cells per plate. The cells were then incubated overnight at 37°C in 5% CO_2_ incubator. The following day the cells were treated with the MAPKi combination, Dabrafenib at a final concentration of 100nM and Trametinib at a final concentration of 10nM. This MAPKi treatment after 48 hours showed a near complete knock out in cell proliferation. These cells were continuously kept in media containing the MAPKi, which was refreshed every 48h. This treatment was continued until resistance was observed in cells. Resistance was defined as the point when cells initiate proliferation in this MAPKi combination. All the cell lines typically took ∼2 months to acquire resistance to the drug treatment. Once cells acquired resistance, they were consistently kept in media containing MAPKi.

### Drug treatments

A375, M481, UACC62, WM164, and 451Lu cells, naïve and MAPKi-resistant, were setup in 24 well plates. These cells were either setup in 2D or 2.5D. For 2.5D assays, collagen was first polymerized in the wells to which the cells were then added. Briefly, cells at ∼70% confluency were trypsinized, harvested, and counted to determine the cell concentration. The cells were then seeded at 10,000 cells/well and incubated overnight to allow attachment to the bottom of the dish. To examine proliferation upon MAPKi treatment, the naïve cells were treated with vehicle (DMSO) control and MAPKi for 48h and compared to resistant cells that are consistently under the pressure of MAPKi. The final DMSO concentration was 0.0001% in vehicle and drug treated cells.

To evaluate the effect and specificity of ROCK inhibition on drug resistant melanoma lines, the naïve and resistant cells were seeded in 24 well plates at 10,000 cells/well and incubated overnight. The cells were then treated with ROCK inhibitors (ROCKi) Y27632 or GSK269962A. The naïve cells only received the ROCKi and the MAPKi-resistant melanoma cells received the MAPKi and the ROCKi. The cells were treated with the ROCKi for 48h.

Effect of PKA activation on MAPKi-resistant melanoma cells was measured by treating cells with a vehicle control and 8-bromo-cAMP for 48h.

### Fluorescent staining for cell fate measurement

Cell fate measurements were performed as described previously (Murali et al., 2019; Siruvallur Murali et al., 2021). Briefly, for cell viability following drug treatment, the media was aspirated from the cells and incubated for 30 minutes with 4µM EtHd and 10µg/mL Hoechst 33342 in fresh phenol red-free DMEM supplemented with 10% FBS and 1% antibiotic-antimycotic. Imaging was then performed with a Nikon *Ti* epifluorescence microscope with an OKO temperature and CO_2_ control system regulated at 37 °C and 5% CO_2_. The filter set for the red channel had an excitation at 540-580 nm and emission at 600-660 nm, and blue channel had an excitation at 340-380 nm and emission at 435-485 nm.

Cell proliferation was quantified by identifying cells undergoing S phase of cell cycle. Briefly, cells were incubated with modified thymidine analogue EdU at a final concentration of 20µM in the treatment media for 1h. The cells in 2D and 2.5D were washed with 1X PBS. The cells cultured in 2.5D were fixed with 4% paraformaldehyde for 30 min at 37 °C, and the cells in 2D for 20 min at room temperature. The fixed cells were permeabilized with 0.5% Triton X-100 for 30 min in 2.5D and 20 min for 2D. The cells were washed with 1X PBS and Click reaction was performed. To make 1mL of Click reaction mix, 888µL of 1X PBS, 100µL of ascorbic acid (200 mg/mL stock), 10µL of copper sulfate (50 mg/mL stock) and 2uL Alexa Fluor Azide-647 (50 µM stock) was used. Cells were then incubated in the reaction mix for 30 mins. (*Note:* reaction mix was added immediately after preparation as the mix begins to lose activity after ∼ 10 to 15 min.) The cells were washed three times with 1X PBS after incubation and then stained with DAPI nuclear stain for 5 min. The cells were then washed and ready for imaging. Imaging was performed on the Nikon *Ti* epifluorescence microscope.

### Cell fate image analysis using cell profiler

The cell fate assays to determine viability and proliferation were imaged at 10X magnification using the Nikon-*Ti* microscope. 2D images were acquired as tiff files and the 2.5D was acquired as a Z-stack following which a maximum intensity Z-projection image was generated. The images were then analyzed using Cell Profiler (http://cellprofiler.org) (Carpenter et al., 2006) as described before (Siruvallur Murali et al., 2021). Briefly, pipelines were developed to classify each channel to segregate for proliferation/viability marker and nuclear marker for total cell number. Illumination correction to remove background signal followed by thresholding was then performed. A random set of images were manually thresholded and this threshold value was then applied to all the images from that experiment. The total number of positive pixels were determined for each channel and ratiometric analysis was done to evaluate viability (total dead pixels/total nuclear pixels) or proliferation (total proliferation positive pixels/total nuclear pixels).

### Glucose uptake assay

For glucose uptake, upon MAPKi treatment in naïve and resistant cells or GSK269962A treatment in resistant cells, the cells were treated with drugs for 36 h in media containing glucose. The media containing glucose was then removed and the cells are treated with drugs in glucose-free media overnight (∼12 h), thus making the total drug treatment to 48 h. NBD-glucose was added to cells in the drug-containing glucose-free media at a final concentration of 50 µg/mL for 3 h. 30 minutes prior to 3 h, Hoechst 33342 was added to NBD-glucose at a final concentration of 10 µg/mL. The cells were then washed twice with PBS and twice with phenol red-free DMEM containing 10% FBS. Fresh phenol red-free media was added and the cells were imaged. Control cells with drug treatments and Hoechst 33342 but without NBD-glucose were prepared to acquire background fluorescence images. The Nikon *Ti*, epifluorescence microscope with a Plan Apo 60X (1.4 NA) oil objective with a lambda-XL light spurce and Lambda 10-B smart shutter were used for image acquisition. Images were acquired with a Zyla CMOS camera (Andor) and the microscope controlled using µManager.

The images were quantified using ImageJ. The intensity for background images where cells were not treated with NBD-glucose but incubated with or without drugs and Hoechst 33342 was quantified as intensity/cell and the average background intensity/cell was then subtracted from each image of cells treated with NBD-glucose. The Hoechst 33342 was used to count the total number of cells. Subsequently intensity/cell for cells with or without drugs but with NBD-glucose and Hoechst33342 that were imaged, quantified and background intensity subtracted to give us intensity/cell upon NBD-glucose treatment.

### Metabolic rate quantification

Seahorse assay to evaluate metabolic rate was performed on the Seahorse XFe24 analyzer (Agilent Technologies) using the Seahorse FluxPaks (102340-100) which contains the Seahorse cartridge (100850-001), Seahorse 24 well plates (100777-004) and XF calibrant (100840-000). The Seahorse XF DMEM assay medium (103680-100) was supplemented with 100mM glucose (103577-100), 5mM pyruvate (103578-100) and 2.5mM glutamine (103579-100). The XF DMEM media along with the supplements was then used for the metabolic rate assays. For seahorse glycolytic rate assays, Rotenone /Antimycin A at a final concentration of 0.5µM and 2-deoxy-D-glucose was used at a final concentration of 50mM. The Rotenon/Antimycin A targets mitochondrial ETC complexes I and III to inhibit mitochondrial respiration which results in increased glycolysis. The 2-deoxy-D-glucose inhibits hexokinase, leading to decreased glycolysis. To perform PKA activation in MAPKi-resistant melanoma cells, 8-bromo-cAMP was purchased from Tocris (Cat No. 1140).

The assay was performed according to manufacturer’s instructions. To evaluate the basal metabolic rate difference between melanoma naïve and resistant cells, the cells were setup at 10,000 cells/well and incubated overnight at 37 °C. The following day the naïve cells were treated with DMSO and MAPKi. The resistant cells were consistently under the presence of MAPKi. The cells were incubated with MAPKi for 48 h. To evaluate the effect of GSK269962A, the cells were treated with GSK269962A at 0, 156.25, 312.5, 625, 1250, 2500 and 5000nM for 48h. 24 h prior to the completion of the assay, a seahorse cartridge was hydrated with the XF calibrant (pH 7.4) and incubated at 37 °C without CO_2_. On the day of the experiment, the seahorse XFe24 analyzer was calibrated using the hydrated cartridge. The cells were then washed once with seahorse XF DMEM and refreshed with 525 µL of fresh seahorse XF DMEM. The cells were then incubated at 37 °C without CO_2_. After incubating the cells for 1 h, the seahorse assay was performed by measuring the extracellular acidification rates as a measure for glycolytic rate and the oxygen consumption rate as a measure for oxidative phosphorylation. On the day of the experiment, the hydrated cartridge was injected with a combination of Rotenone plus Antimycin A (Rot/AA) in port A at a final concentration of 0.5µM and 2-deoxy-D-glucose (2-DG) in port B at a final concentration of 50mM.The seahorse XFe24 analyzer was then calibrated using the hydrated cartridge and then the glycolytic rate assay was then performed on the plate with cells.

To normalize the seahorse results to cell number, after completion of the assay the cells in the dish were fixed with 4% paraformaldehyde and DAPI stained. The cells were then imaged on Nikon-*Ti* inverted epifluorescence microscope with Plan Fluor 10X (?? NA) objective and a motorized stage with X-cite 120 LED light source (Lumen Dynamics). Images were acquired as 8 X 8 stitched fields with 10% overlap to fully capture the area of a single well using a Zyla CMOS camera (Andor) controlled by Nikon Elements Software.

### Metabolomics

A375 naïve and resistant cells were seeded at a density of 1 x 10^6^ cells per dish. The following day, the naïve cells were treated with DMSO or MAPKi. The resistant cells were under the pressure of MAPKi and incubated for 48 h. The cells were scraped and collected after they were washed thrice with ice-cold normal saline and lysed with ice-cold 80% methanol in water. All lysed samples were scraped into Eppendorf tubes followed by three freeze-thaw cycles in liquid nitrogen and then centrifuged and pelleted at 4 °C. The supernatant from this centrifugation was then collected and dried in an Eppendorf tube using a SpeedVac concentrator (Savant, Thermo Fisher Scientific). The metabolites were rehydrated in 100uL of 0.03% formic acid (85178, Thermo Fisher Scientific) in liquid chromatography-mass spectrometry (LC-MS)-grade water (51140, Thermo Fisher Scientific), vortex-mixed to remove debris, and centrifuged. The supernatant was transferred to a high-performance liquid chromatography (HPLC) vial, and metabolite profiling was determined by liquid chromatography-tandem mass spectrometry (LC-MS/MS). An AB QTRAP 5500 liquid chromatography-triple quadrupole mass spectrometer (Applied Biosystems SCIEX) was used for chromatogram review and peak area integration. The peak area for each metabolite was normalized to the total ion count of that sample (Supplementary Table ???), and the normalized data were auto-scaled (mean centered and divided by the s.d. of each variable) using MetaboAnalyst 5.0 (www.metaboanalyst.ca) (Chong et al., 2018). Univariate statistical differences of metabolites between the groups were analyzed using a two-tailed Student’s *t*-test.

### Western Blots and Activity Assays

Cells were washed twice in chilled 1X PBS and lysed in chilled 1X RIPA buffer supplemented with protease (Roche, Cat No. 04 693 116 001) and phosphatase inhibitors (Roche, Cat No. 04 906 837 001). Protein concentrations were determined using Pierce BCA assay kit. Samples were run on 4-20% Mini-PROTEAN pre-cast gels (Biorad, Cat No. 4568094) and transferred to PVDF membranes. The RhoActivity assay was performed using the RhoA pull-down activation assay biochem kit (Bead pull-down format) (Cytoskeleton, Cat No. BK036). The assay was carried out by incubating 800 µg of total cell lysate with 50 µg of Rhotekin as described in the manufacturers protocol following which western blots were performed. Antibodies used and dilutions are as follows: RhoA 1:1000 (Cell Signaling Technology, 2117), β-actin 1:5000 (Cell Signaling Technology, 3700S), Myosin Light Chain 2 1:1000(Cell Signaling Technology, 3672), Thr18/Ser19 Phospho-Myosin Light Chain 2 1:1000(Cell Signaling Technology, 3674), GLUT1 1:1000 (Cell Signaling Technology,12939S), GLUT4 1:1000 (Abcam, ab33780), Insulin receptor substrate 1 1:1000(Cell Signaling Technology, 2383L), Ser636/639 Phospho-Insulin receptor substrate 1 (Cell Signaling Technology, 2388S). The western blot gels were quantified by evaluating the band intensities using ImageJ (Wang et al., 2017).

### Immunofluorescence

Immunofluorescence was performed on cells grown on no. 1.5 glass coverslips that had been previously sterilized in ethanol and washed twice with 1X PBS. The cells after seeding on the coverslip were treated with inhibitors for 48h and then fixed with pre-warmed 4% paraformaldehyde in PBS for 10 min at 37 °C. The cells were then washed and permeabilized with 0.5% Triton X-100 for 20 mins. The coverslips were then blocked with 5% BSA in 1X PBS for 1 h followed by incubation of the primary antibody overnight at 4°C in a humidified chamber. The cells were then washed and treated with Alexa-fluor conjugated secondary antibody for 1h followed by washing. To stain the actin structures, the cells were incubated with Alexafluor-conjugated phalloidin for 1 h and finally stained with DAPI for 10 mins. The coverslips were washed and mounted with Fluormount mounting medium for microscopy. The fixed cells were then imaged on a Nikon-*Ti* inverted ffluorescence microscope with a Plan Apo 60X (1.4 NA) oil objective with a lambda-XL light spurce and Lambda 10-B smart shutter. Images were acquired with a ZylaCMOS camera (Andor) and the microscope controlled using µManager. Antibodies used and the dilutions are as below: Phalloidin 1:50 from a 40X stock solution (Invitrogen, A12379); 2-NBD glucose at a final concentration of 100µg/mL (Cayman, 11046); Anti-phosphotyrosine antibody, clone 4G10 1:50 (Millipore Sigma, 05-321).

### Protein kinase A (PKA) Activation Assay

A colorimetric assay was used to quantitatively evaluate the activation of PKA. This assay uses an immobilized PKA substrate that is bound to the microtiter plate. The samples containing PKA in the presence of ATP phosphorylate the immobilized PKA substrate upon incubation. An antibody specific to phospho-PKA substrate binds to the modified immobilized substrate. Substrate is then added, and the intensity of color developed is directly proportional to the amount of active PKA in the sample. The kit was used as per the manufacturers protocol. Briefly, the cells were lysed in lysis buffer containing 1mM phenylmethylsulfonyl fluoride (PMSF), 10mM sodium orthovanadate, and centrifuged to collect the supernatant. BCA assay was performed to evaluate protein concentrations. The standards and samples were diluted in 1X kinase reaction buffer. The strips were then prepared to which 40µL of standard or diluted sample, 10µL of reconstituted ATP were added and incubated for 90 minutes at 30°C. The wells were then washed and added with 25µL Donkey anti-rabbit IgG HRP and 25µL of Phospho PKA substrate antibody and incubated for 60 mins. The wells were then washed, 100µL of TMB substrate chromogen was added and incubated for 30 mins followed by 50µL of stop solution. The absorbance was read at 450nm, and the results plotted with the standard curve.

### Flow cytometry analysis

To evaluate glucose transporters on cells, the cells were harvested with TryLE, washed with cold PBS and centrifuged and counted. 1 X 10^6^ cells were collected for each condition (a total of four conditions) in 1.5mL Eppendorf tube and kept steps performed on ice. Each tube was treated with either anti GLUT1, anti GLUT3 or anti GLUT4 antibody and one kept for negatives. The cells were incubated with antibody at a final dilution of 1:50 for 30 minutes on ice and mixed every 10 minutes. These cells were then washed with cold PBS and centrifuged and subsequently fixed for 15 minutes on ice. After fixation the cells were washed with cold PBS and treated with Alexa fluor 488 for GLUT1, Alexa Fluor 568 for GLUT3 and Alexa Fluor 647 for GLUT4 on ice for 30 minutes and washed with cold PBS. These cells were then passed through a cell filter to acquire single cells and flow cytometry was then performed. The results to determine percent positive cells was done on FlowJo by gating the negative cells and then applied to all stained cells.

### Plasmids and virus packaging

Plasmid expressing constitutively activate RhoA was ordered from Addgene (pSLIK CA RhoA, Plasmid #112894) (MacKay and Kumar, 2014). Since the pSLIK had a Venus tag, the CA RhoA was cloned into a pLVX-cmv100-puro lentiviral vector. For virus packaging of the pSLIK lentiviral construct, psPAX2 and pMD2.G were used (Addgene # 12260 and Addgene # 12259); plasmids were a gift from D. For lentivirus production, HEK293 cells were transfected with expression and packaging plasmids following standard calcium phosphate or polyethylenimine (Polyscience, 23966) protocols. The supernatant was collected after two days transfection, filtered through 0.45µm mixed cellulose esters membrane syringe filters (Fisher Scientific, Cat no. 09-720-005) and incubated on A375 naïve cells in the presence of 2µg/mL polybrene (Millipore, Cat no. TR-1003-G). The cells were then selected and used for a maximum of 3 passages post-selection. The expression of RhoA was validated through myosin and phosphorylated myosin light chain expression (pMLC).

### Statistical Analysis

Statistical analysis and data plots were prepared using Graphpad Prism 9.5.1 as described in the figure legends. The biological and technical replicates for each experiment have been described in the figure legends. The statistical significance for all the results is: *P<0.05; **P<0.01; ***P<0.001; ****P<0.0001.

