## Supplementary figures and images for "RhoA activation promotes glucose uptake to elevate proliferation in MAPK inhibitor resistant melanoma cells"

### Supplemental Figure 1

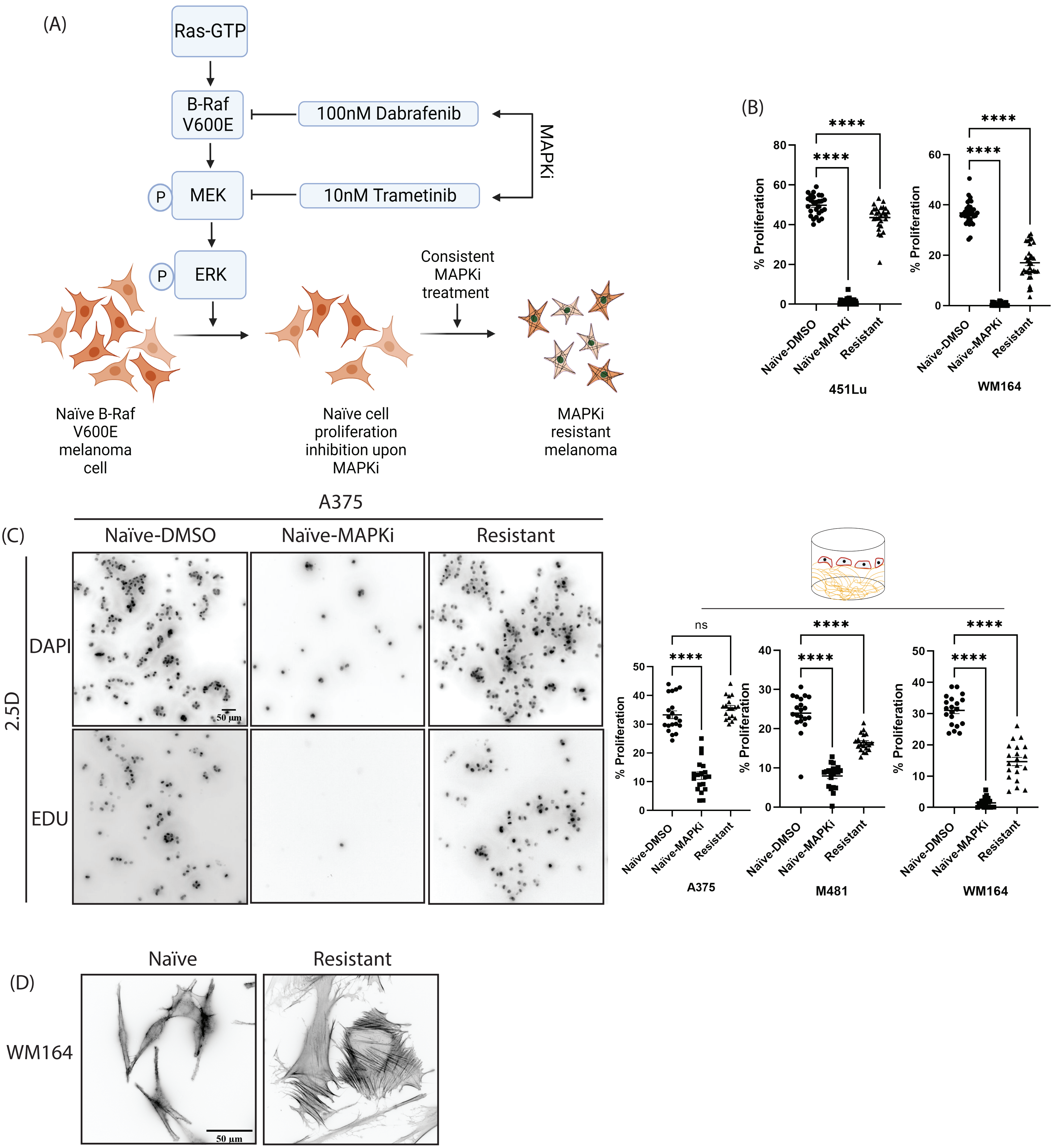

### Supplemental Figure 2

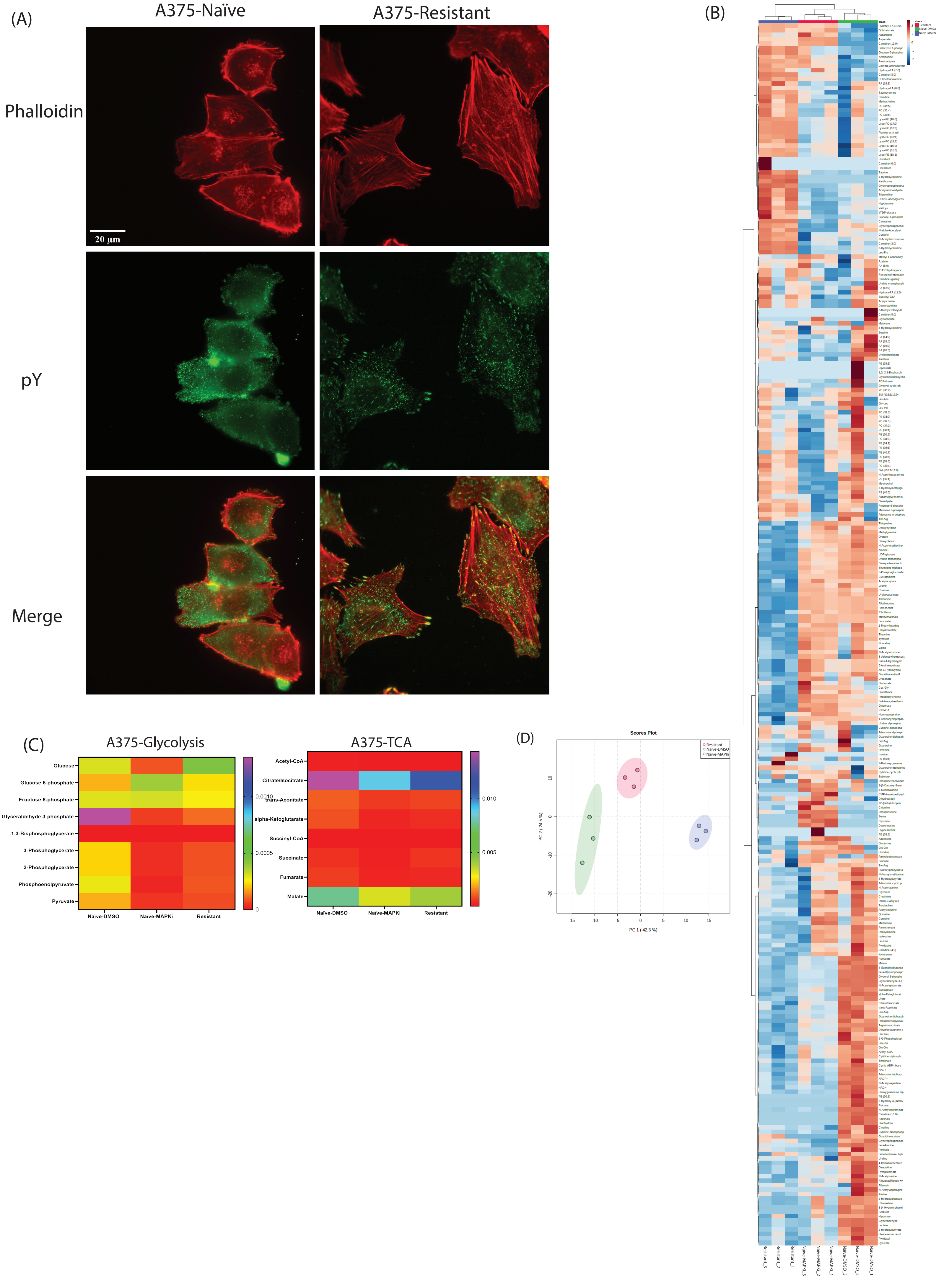

### Supplemental Figure 3

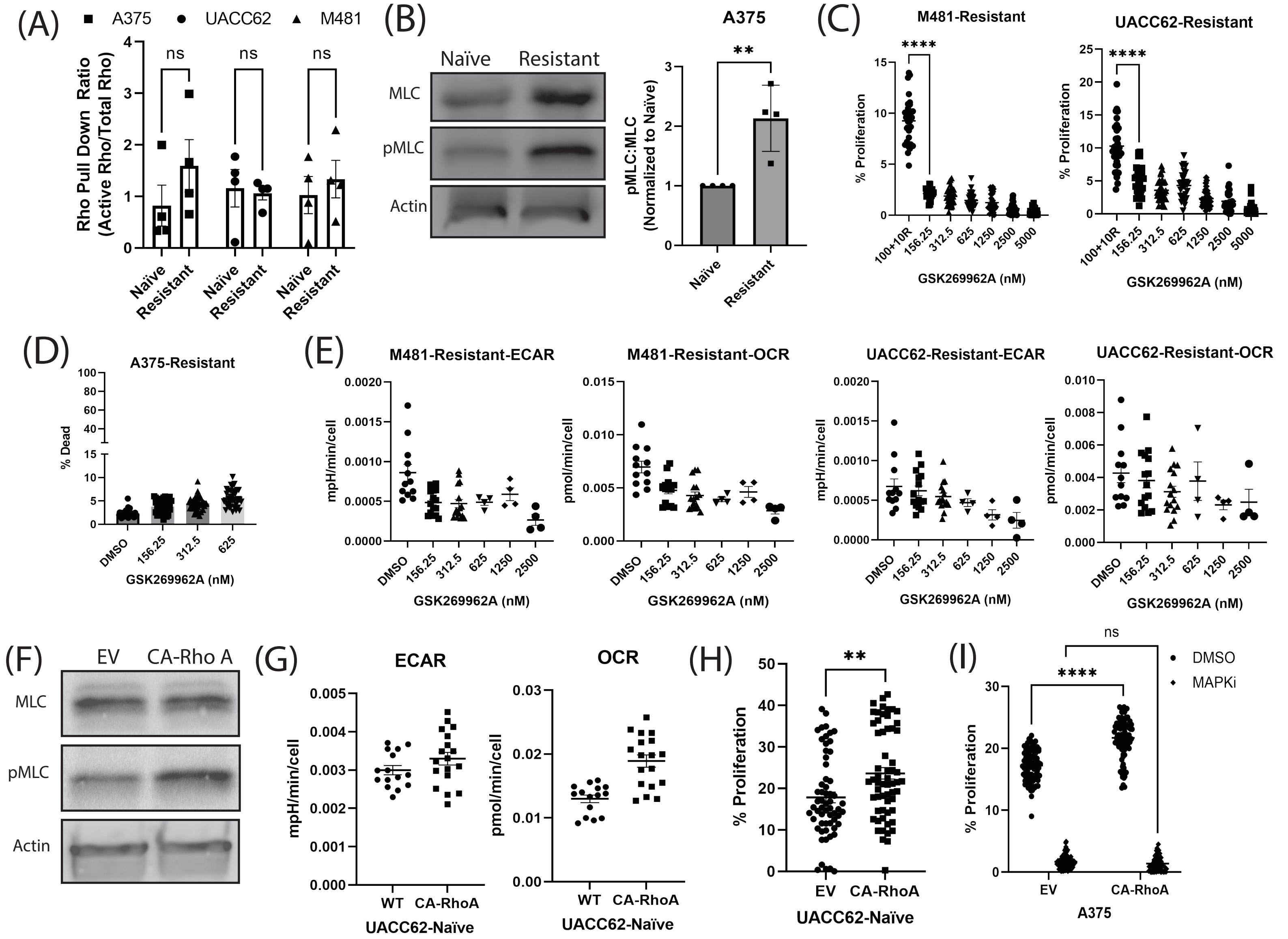
