## Supplemental Figure 4 for "RhoA activation promotes glucose uptake to elevate proliferation in MAPK inhibitor resistant melanoma cells"

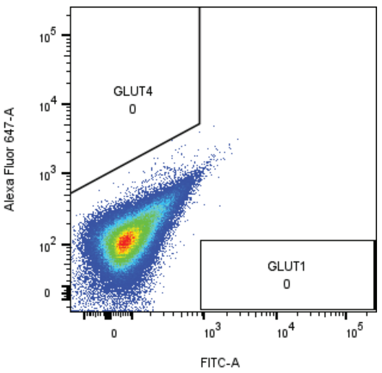


A375-Naïve-Negative

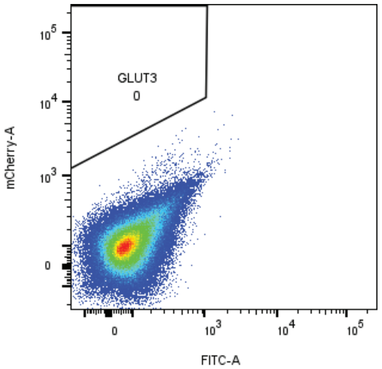


A375-Naïve-Negative

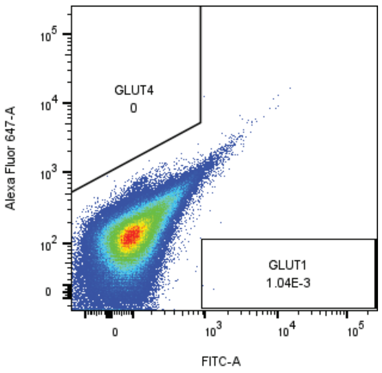


A375-Resistant-Negative

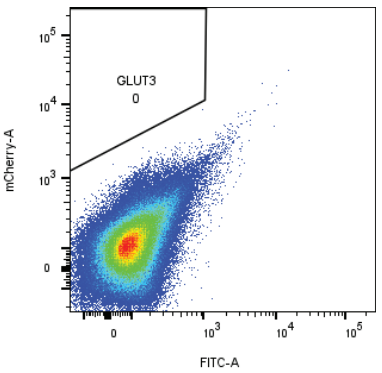


A375-Resistant-Negative

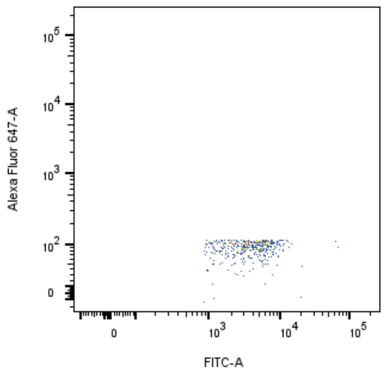


A375-Naïve-GLUT1

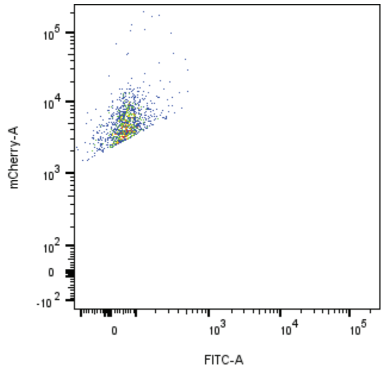


A375-Naïve-GLUT3

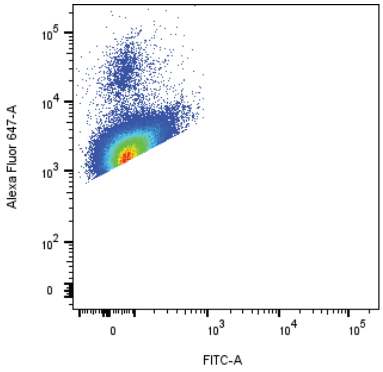


A375-Naïve-GLUT4

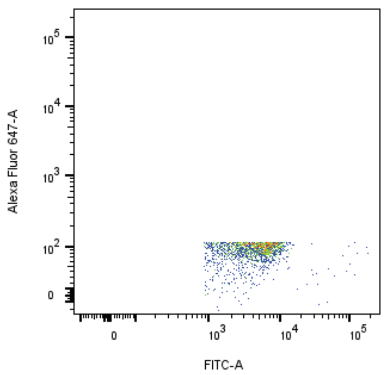


A375-Resistant-DMSO-GLUT1

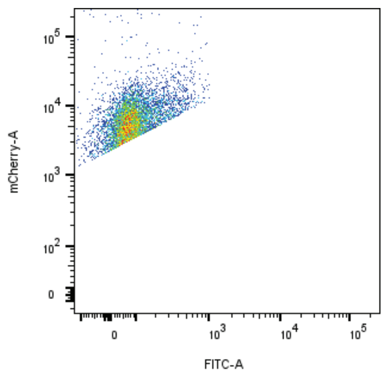


A375-Resistant-DMSO-GLUT3

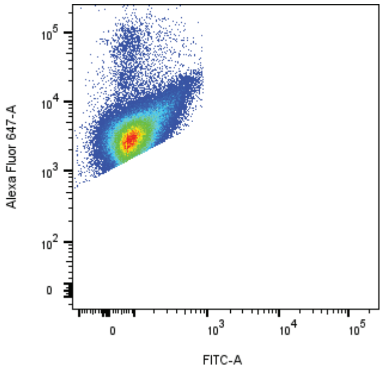


A375-Resistant-DMSO-GLUT4
